# Comparative Genomics of Six *Juglans* Species Reveals Disease-associated Gene Family Contractions

**DOI:** 10.1101/561738

**Authors:** Alexander Trouern-Trend, Taylor Falk, Sumaira Zaman, Madison Caballero, David B. Neale, Charles H. Langley, Abhaya Dandekar, Kristian A. Stevens, Jill L. Wegrzyn

## Abstract

*Juglans* (walnuts), the most speciose genus in the walnut family (Juglandaceae) represents most of the family’s commercially valuable fruit and wood-producing trees. It includes several species used as rootstock in agriculture for their resistance to various abiotic and biotic stressors. We present the full structural and functional genome annotations of six *Juglans* species and one outgroup within Juglandaceae (*Juglans regia, J. cathayensis, J. hindsii, J. microcarpa, J. nigra, J. sigillata* and *Pterocarya stenoptera*) produced using BRAKER2 semi-unsupervised gene prediction pipeline and additional tools. For each annotation, gene predictors were trained using 19 tissue-specific *J. regia* transcriptomes aligned to the genomes. Additional functional evidence and filters were applied to multi-exonic and mono-exonic putative genes to yield between 27,000 and 44,000 high-confidence gene models per species. Comparison of gene models to the BUSCO embryophyta dataset suggested that, on average, genome annotation completeness was 85.6%. We utilized these high-quality annotations to assess gene family evolution within *Juglans* and among *Juglans* and selected Eurosid species. We found notable contractions in several gene families in *J. hindsii*, including disease resistance-related Wall-associated Kinase (WAK) and *Catharanthus roseus* Receptor-like Kinase (CrRLK1L) and others involved in abiotic stress response. Finally, we confirmed an ancient whole genome duplication that took place in a common ancestor of Juglandaceae using site substitution comparative analysis.

## INTRODUCTION

It is anticipated that new genomic resources for *Juglans* (walnuts) will lead to improved timber and nut production by accelerating development of advanced agriculture, breeding efforts, and resource management techniques for the genus. Already, these practices are beneficiaries of the genomic analyses and tool development potentiated by the growing pool of *Juglans* sequence data (Martínez-García *et al*. 2017; Bernard *et al*. 2018; Marrano *et al*. 2018; Famula *et al*. 2019; Zhu *et al*. 2019). The recent publication of the unannotated draft reference genomes of six *Juglans* species: *J. nigra* (Eastern black walnut), *J. hindsii* (Hinds black walnut), *J. microcarpa* (Texas walnut), *J. sigillata* (iron walnut), *J. cathayensis* (Chinese walnut), *J. regia* (Persian or English walnut) and a member of the sister group to *Juglans, Pterocarya stenoptera* (Chinese wingnut) greatly expands the existing resource and provides an unprecedented opportunity to apply tools such as genomic selection to the Juglandaceae (Stevens *et al*. 2018).

The genomes considered are Eurasian and North American species from across the three sections of the genus, *Cardiocaryon, Dioscaryon* (syn. sect. *Juglans*) and *Rhysocaryon* (Figure 1). These species were selected for their historical and agricultural importance and for their phylogenetic placements, which span the breadth of the genus. The North American species *J. nigra, J. hindsii*, and *J. microcarpa* are members of sect. *Rhysocaryon* and grow in riparian forests in the eastern, western and southern United States, respectively. *J. hindsii* and *J. microcarpa* are distributed in moderately dispersed populations. Conversely, *J. nigra*’s range is more contiguous and expansive and overlaps with the northeastern distribution of *J. microcarpa*. Among these, *J. nigra* is especially valued for its high-quality timber, cold-hardiness, and disease resistance (Beineke 1983; Settle *et al*. 2015; Chakraborty *et al*. 2015; Chakraborty *et al*. 2016). *J. hindsii* is characterized as vigorous, drought tolerant, and resistant to *Armillaria* root rot (honey fungus) (Buzo *et al*. 2009). The need for disease-resistant and climate-tolerant cultivars for nut production has driven the use of *J. nigra, J. hindsii* and *J. microcarpa* in hybridization trials (Browne *et al*. 2015). These species contribute resistance to many soilborne pathogens, including *Phytophthora* and a range of nematodes.

**Figure 1:**
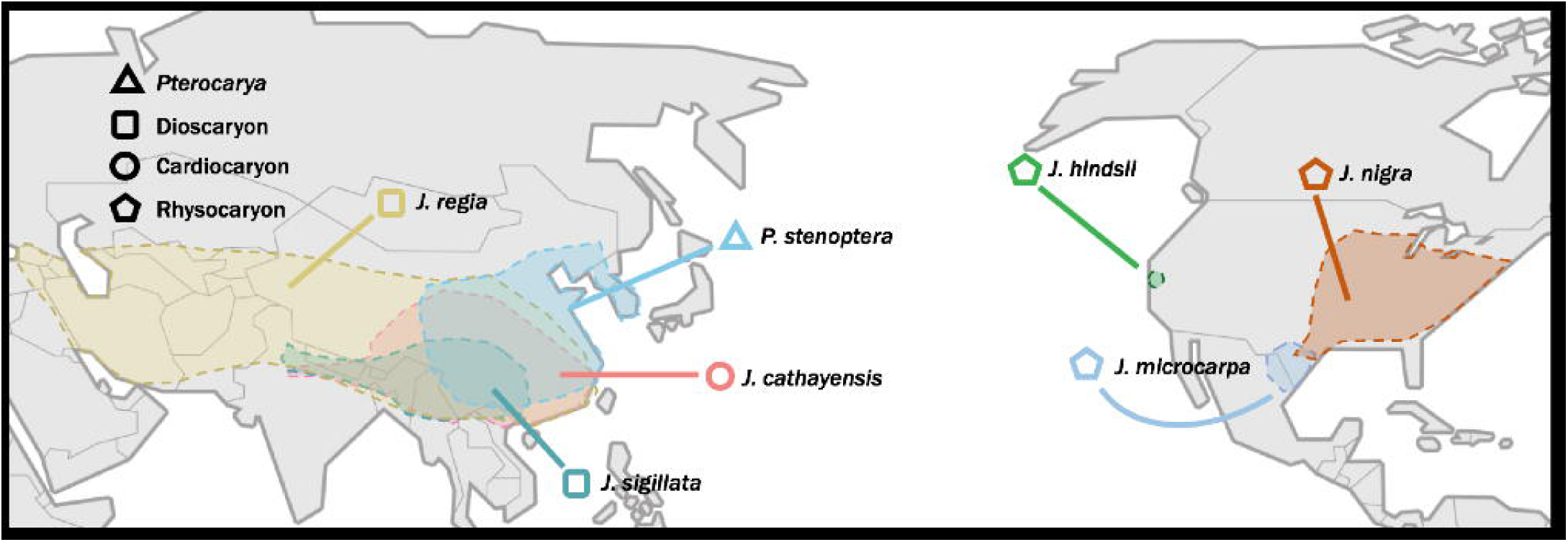
Approximate ranges of each annotated species with shapes denoting the three sections of *Juglans* and the outgroup genus, *Pterocarya*.

The sampled Eurasian species include *J. regia*, the predominant nut producer of all cultivated *Juglans. J. regia*’s western native range continues into Southeastern Europe, an impressive relic of silk road trading that underlines its agricultural suitability (Pollegioni *et al*. 2015). *J. sigillata* is phylogenetically adjacent to *J. regia* as the only other species of sect. *Dioscaryon*, and, like *J. regia*, is valued for both its timber and nut (Weckerle *et al*. 2005). Despite their divergent morphology, which includes number of leaflets and nut characteristics, a growing body of molecular evidence suggests that *J. regia* and *J. sigillata* may not qualify as separate species (Gunn *et al*. 2010; Zhao *et al*. 2018). *J. sigillata* grows sympatrically with *J. regia* and *J. cathayensis* (sect. *Cardiocaryon*) in southwestern China, but the distribution of *J. cathayensis* extends beyond the sympatric zone several hundred kilometers eastward to the coastline and northward towards the Gobi desert. *J. cathayensis* is endangered in its natural range in China (Zhang *et al*. 2015) and is evaluated in breeding programs for its resistance to lesion nematodes. The outgroup, *P. stenoptera* is also native to southeastern China and is used for ornamental planting, timber, and medicinal extracts. *P. stenoptera* has been integrated into hybridization trials as a non-viable (inconsistent grafting) rootstock for its resistance to *Phytophthora* (Browne *et al*. 2011).

Stevens *et. al* (2018) annotated a set of microsyntenic regions containing polyphenol oxidase loci to confirm a gene duplication in an ancestral *Juglans*. Here, we describe the full gene annotations of these diploid genomes, a critical missing component to their full utilization as a genomic resource. We demonstrate their utility by investigating the genomes in a comparative manner. First, we view the evolution of gene families in *Juglans* across the phylogeny. Second, we leverage the annotations for a comparative genomic analysis to date an ancient whole genome duplication in an ancestral species of Juglandaceae.

## RESULTS

### Semi-unsupervised Gene Prediction

Errors introduced from genome annotation often lead to inconsistent gene expression estimates and contribute to the inaccurate characterization of gene space, gene family evolution and timing of whole genome duplications (Vijay *et al*. 2013, Denton *et al*. 2014). Our approach was applied across all seven genomes that leveraged RNA-Seq reads generated from tissue-specific libraries of *J. regia* (Table 1). This approach took advantage of the deep sequencing by directly aligning reads to the genome to resolve challenges associated with reliance on error-prone and often fragmented *de novo* assembled transcripts (Hoff *et al*. 2016). The bias introduced by using RNA-Seq reads solely from *J. regia* for the annotation of all genomes was partially mitigated by the semi-supervised training of the gene prediction tool, AUGUSTUS, included in BRAKER. The AUGUSTUS component utilizes the evidence of successful alignments to learn features of the genome in question and propose gene models. Repeat libraries were generated and subsequently used for masking between roughly 44% and 48% of the genomes prior to read alignment (Table 2; Table S1). The raw reads aligned across the genomes at rates inversely proportional to their phylogenetic distance from *J. regia*. Average alignment rates across *J. regia* transcriptome libraries are displayed as total mapped (concordantly mapped): 87.1% (84.2%) in *J. regia*, 88.3% (78.7%) in *J. sigillata*, 51.8% (49.1%) in *J. cathayensis*, 68.0% (49.0%) in *J. nigra*, 41.0% (38.7%) in *J. microcarpa*, 49.7% (48.0%) in *J. hindsii* and 33.2% (31.2%) in the outgroup, *P. stenoptera*. Initial gene prediction estimates from BRAKER2 ranged from 81,753 (*J. hindsii*) to 133,963 (*P. stenoptera*) (Table S2). Filtering of BRAKER2 models considered completeness (start and stop codon present), isoforms, exon lengths, intron lengths, and splice sites. These considerations provided a reduced set of gene models for each genome: 82,610 in *J. nigra*, 76,847 in *J. hindsii*, 114,573 in *J. microcarpa*, 83,457 in *J. sigillata*, 97,312 in *J. cathayensis*, 84,098 in *J. regia*, and 123,420 in *P. stenoptera* (Table S2).

**Table 1:** Genome assembly statistics for seven Juglandaceae species

|  | *Juglans.*<br>*hindsii* | *Juglans*<br>*nigra* | *Juglans*<br>*cathayensis* | *Juglans*<br>*microcarpa* | *Juglans*<br>*sigillata* | *Juglans*<br>*regia* | *Pterocarya*<br>*stenoptera* |
| --- | --- | --- | --- | --- | --- | --- | --- |
| Plant Info |  |  |  |  |  |  |  |
| Name | ‘Rawlins’ | ‘Sparrow’ | ‘Wild Walnut’ | ‘83-129’ | ‘Yangbi 1’ | ‘Chandler’ | ‘83-13’ |
| Cultivar | DJUG105 | A30 | DJUG11.03 | DJUG29.11 | DJUG951.04 | 64-172 | DPTE1.09 |
| Source | NCGR | MU | NCGR | NCGR | NCGR | UCD | NCGR |
| Assembly |  |  |  |  |  |  |  |
| Version | 1.0 | 1.0 | 1.0 | 1.0 | 1.0 | 1.1 | 1.0 |
| Size (Mbp) | 605.70 | 605.05 | 751.37 | 896.45 | 622.24 | 686.52 | 936.89 |
| Scaffolds | 273,094 | 232,579 | 332,634 | 329,873 | 282,224 | 186,636 | 396,056 |
| N50 (Kbp) | 512.79 | 118.45 | 158.25 | 145.01 | 218.35 | 278.29 | 159.70 |
| Annotated Assembly (> 3Kbp scaffolds) |  |  |  |  |  |  |  |
| Size (Mbp) | 586.05 | 580.70 | 719.60 | 862.79 | 585.63 | 686.52 | 902.23 |
| Scaffolds | 4,672 | 5,896 | 10,342 | 12,024 | 6,413 | 11,848 | 11,574 |
| N50 (Kbp) | 540.03 | 271.37 | 168.53 | 151.91 | 238.36 | 278.30 | 167.10 |

**Table 2:** Structural and functional annotations for seven Juglandaceae species

|  | *Juglans.*<br>*hindsii* | *Juglans*<br>*nigra* | *Juglans*<br>*cathayensis* | *Juglans*<br>*microcarpa* | *Juglans*<br>*sigillata* | *Juglans regia* | *Pterocarya*<br>*stenoptera* |
| --- | --- | --- | --- | --- | --- | --- | --- |
| Structural Annotation |  |  |  |  |  |  |  |
| Repeat Content | 46.97% | 47.34% | 48.03% | 46.87% | 46.69% | 47.96% | 43.89% |
| Total Genes | 28,664 | 28,335 | 34,066 | 41,611 | 26,835 | 30,626 | 44,318 |
| Total Complete, Multi-exonics | 24,500 | 23,290 | 28,915 | 35,319 | 22,898 | 26,166 | 36,984 |
| Total Complete, mono-exons | 4,164 | 4,426 | 5,151 | 6,292 | 3,937 | 4,460 | 7,334 |
| Gene Length (Avg) | 4,406.38 | 4,301.45 | 4,193.80 | 4,008.01 | 4,373.40 | 4,235.02 | 3,944.81 |
| CDS Length (Avg) | 1,267.37 | 1,277.62 | 1,220.28 | 1,199.12 | 1,250.97 | 1,220.42 | 1203.44 |
| Exons per Gene (Average) | 6.30 | 6.38 | 6.06 | 5.98 | 6.32 | 6.14 | 5.91 |
| Canonical Splice Sites (%) | 98.70% | 98.67% | 98.69% | 98.70% | 98.60% | 98.77% | 98.76% |
| **Functional Annotation** |  |  |  |  |  |  |  |
| EnTAP (Similarity Search) | 23,822 | 23,607 | 27,815 | 33,711 | 22,036 | 25,420 | 35,771 |
| EnTAP (Gene Family only) | 4,842 | 4,728 | 6,251 | 7,900 | 4,799 | 5,206 | 8,547 |

### Functional Annotation Filtering of Gene Models

Functional annotation via sequence similarity search (SSS) and gene family assignment (GFA) provides a form of validation for the proposed models and assesses their completeness. The reciprocal style search required query and target sequence coverage to pass a set threshold, which helped to eliminate unlikely models and validate the final models. A total of 29,046 models with both SSS result and GFA, and 5,190 with only GFA composed the final *J. nigra* set. The same approach was used for the 29,382 models in *J. hindsii*, 43,051 in *J. microcarpa*, 27,596 in *J. sigillata*, 34,857 in *J. cathayensis*, 31,621 in *J. regia*, and 45,808 in *P. stenoptera* (Table 2; Table S2). Structural assessment of the genes examined splice sites, exons per gene, CDS lengths, and intron lengths (Table 2) and reported average gene/CDS lengths relative to other angiosperm species. The vast majority (> 98% in all species) of the splice sites were canonical (GT/AG). All other splice site detected were GC/AG variants.

### Benchmarking Genome Annotation Completeness

The embryophyta collection of 1440 single copy orthologs derived from OrthoDB can be accessed via BUSCO to estimate the completeness of a plant genome assembly, transcriptome, or set of gene models. These 1440 genes (Embryophyta *odb9*) were aligned to all Juglandaceae assemblies and final gene models (Figure 2; Table S3, Table S4). Across members of *Juglans* genus, BUSCO identified 87 to 93% of their database when evaluated against the genome (Table S5). When provided with filtered complete (full-length) gene models, BUSCO reported 77 to 86% completeness, and 78 to 89% with partial (5’ or 3’ complete) models (Table S4).

**Figure 2:**
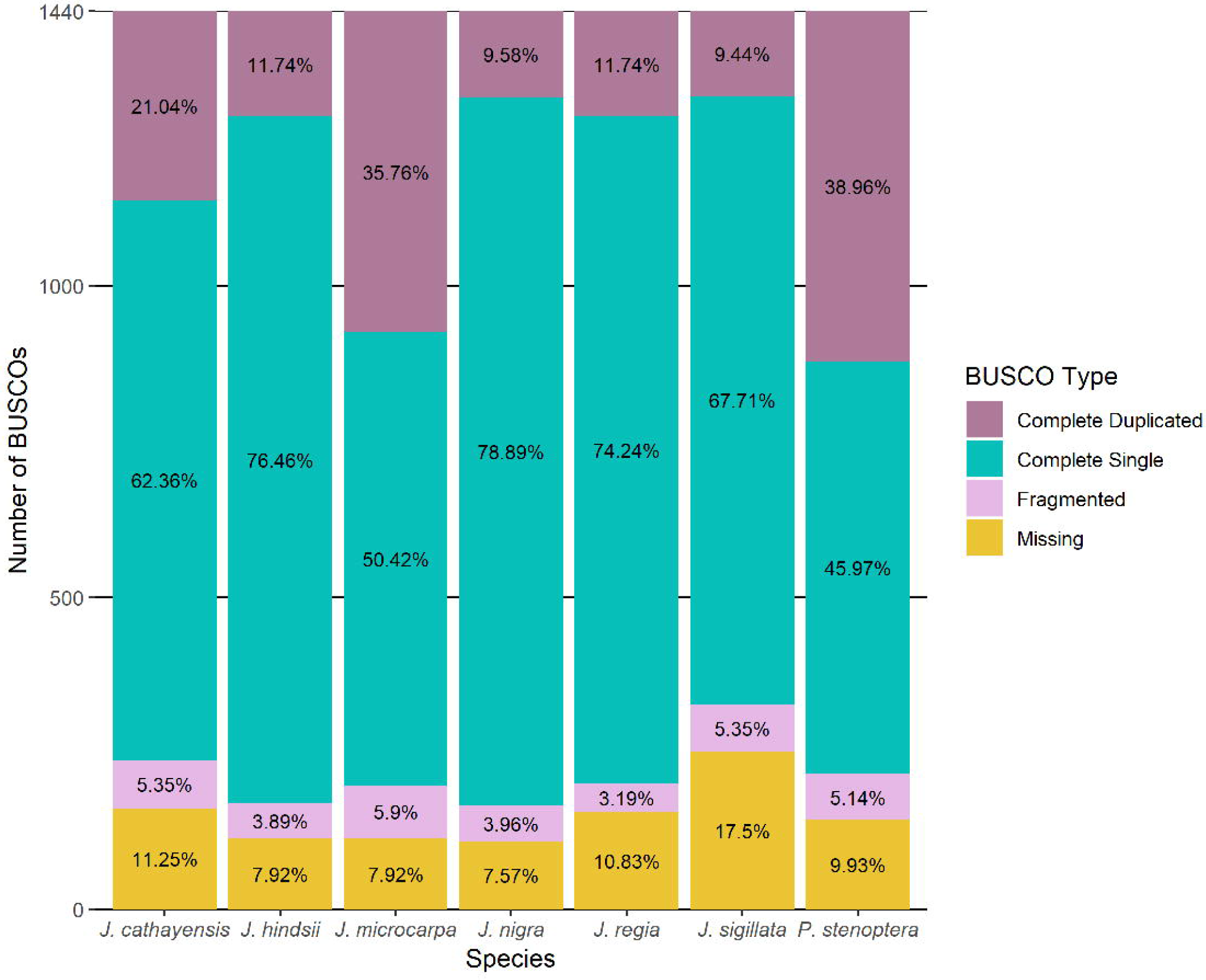
As a measure of gene annotation completeness, the gene models from the seven annotations were compared to a set of 1,440 embryophyta putative single-copy orthologs using BUSCO (Benchmarking Universal Single-copy Orthologs) (Table S3). Orthologs were either found once in the gene models (Complete Single), multiple times (Complete Duplicated), partially (Fragmented), or were not found at all (Missing).

### Orthologous Group Construction

Two OrthoFinder analysis that differed in adjacent clade inclusivity were used to identify homology relationships between genes of the selected species. One run of annotated Juglandaceae (6 *Juglans*, 1 *Pterocarya*) species, and another including previously annotated genomes from across the Eurosid superorder (13 species, see Methods). OrthoFinder assigned 216,778 (92.5%) of the 234,455 total genes from the Juglandaceae set to 26,458 orthogroups (Figure 3B, File S1). The resulting orthogroups range in size from 2 to 190 genes. A total of 161 genes (0.1%) are in 56 species-specific orthogroups. Of the *Juglans* species, *J. cathayensis* had the most genes designated to species-specific orthogroups (24 genes in 8 orthogroups). Just over half, 14,429 orthogroups, have gene membership from all species. A total of 661 orthogroups (5268 genes) are represented by all *Juglans* species (excluding *Pterocarya*). The Juglandaceae set included 538 orthogroups (1159 genes) specific to *Juglans* sect. *Dioscaryon* and 437 orthogroups (1608 genes) specific to members of *Juglans* sect. *Rhysocaryon*. Within *Rhysocaryon*, 905 genes formed 389 orthogroups specific to the parapatric species, *J. microcarpa* and *J. nigra*, but not found in the geographically isolated *J. hindsii*. A total of 149 orthogroups (564 genes) were specific to the three Eurasian *Juglans* species (*J. regia, J. sigillata* and *J. cathayensis*). Genes that could not be assigned to orthogroups, included: 2025 (6.6%) in *J. regia*, 2801 (8.2%) in *J. cathayensis*, 1138 (4.0%) in *J. hindsii*, 3181 (7.6%) in *J. microcarpa*, 934 (3.3%) in *J. nigra*, 1251 (4.7%) in *J. sigillata*, and 6347 (14.3%) in *P. stenoptera* (Figure 3A; Figure 3B; File S1).

**Figure 3:**
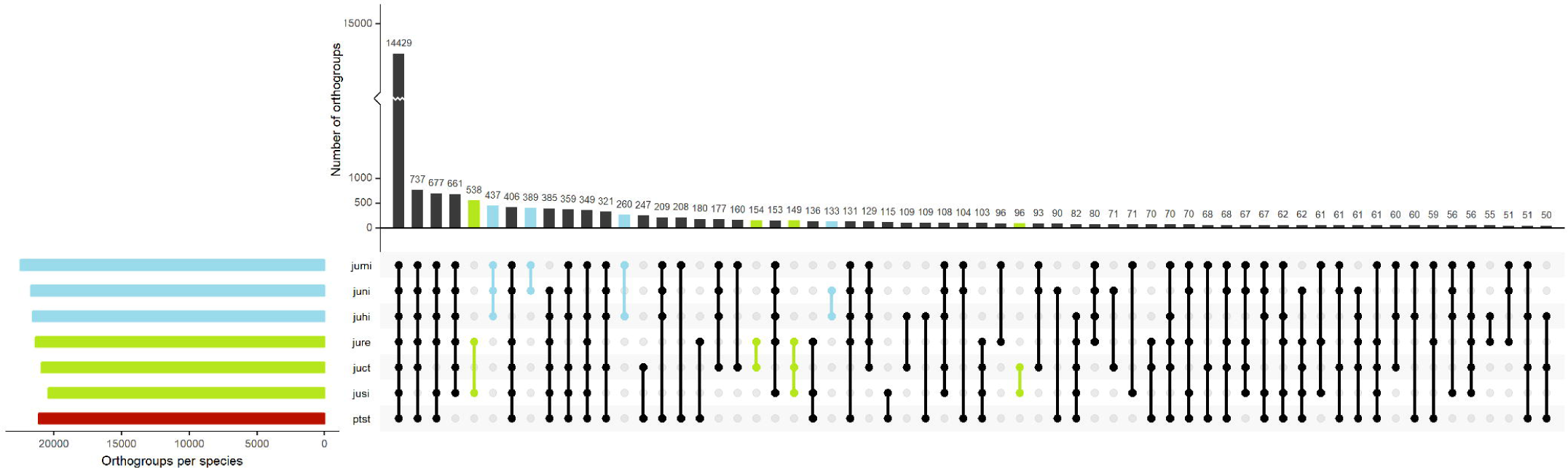
(A) Distribution of species membership across orthogroups. Tiling beneath the histogram indicates the species contributing gene models to each orthogroup in the set. Set size is displayed as height on histogram. The horizontal histogram indicates the number of orthogroups found in each species. Blue indicates data from groups composed of *Rhysocaryon* species while green bars show Eurasian species (*Dioscaryon* and *Cardiocaryon*). B) Cladogram with associated stacked histogram reflecting the number of genes belonging to orthogroups specific to the color-indicated groups.

OrthoFinder analysis of selected Eurosid species assigned 401,186 (92.8.%) of the 456,424 total genes to 22,189 orthogroups (File S2). Of these, a total of 3054 genes (0.7%) are present in 488 species-specific orthogroups and 6722 orthogroups contained at least one gene from each species. The addition of peripheral species to the analysis resulted in an increased gene contribution per species in the orthogroups. This trend is reflected by fewer orthogroups resulting from the Eurosid clustering and the approximate halving of the number of unassigned *Juglans* genes in the Eurosid clustering when compared to the Juglandaceae clustering (Table S6).

### Analysis of Gene Family Evolution

An evaluation of gene families among the annotated species was successful in detecting significant changes between taxa. Prior to gene family analysis with CAFE, orthogroups were filtered to exclude large families (> 100 gene copies) and those composed entirely of paralogs. This removed 57 of 26,458 (0.2%) orthogroups from the Juglandaceae set, and 1880 of 22,189 (8.5%) orthogroups from the Eurosid set. Calculated lambda values were 0.02396 and 0.02197 for Juglandaceae and Eurosid sets, respectively. The higher lambda of Juglandaceae set indicates a higher calculated average rate of gene family evolution. Of the 460 significant rapidly evolving orthogroups discovered based on the Eurosid set, 153 (+131 families expanded/-22 families contracted) had significant changes in *J. microcarpa*, 102 (+57/-45) in *J. regia*, 86 (+62/-24) in *J. cathayensis*, 76 (+30/-46) in *J. sigillata*, 61 (+22/-39) in *J. nigra*, 58 (+32/-26) in *J. hindsii*, and 139 (+113/-24) in *P. stenoptera* (Figure 5, File S4). The Juglandaceae set revealed 430 significant rapidly evolving gene families of which 168 (+123/-45) had significant size changes in the *J. microcarpa* terminal branch, 141 (+86/-55) in *J. regia*, 92 (+72/-20) in *J. cathayensis*, 98 (+39/-59) in *J. sigillata*, 101 (+32/-69) in *J. nigra*, 77 (+40/-37) in *J. hindsii*, and 98 (+67/-31) in *P. stenoptera* (File S3).

**Figure 5:**
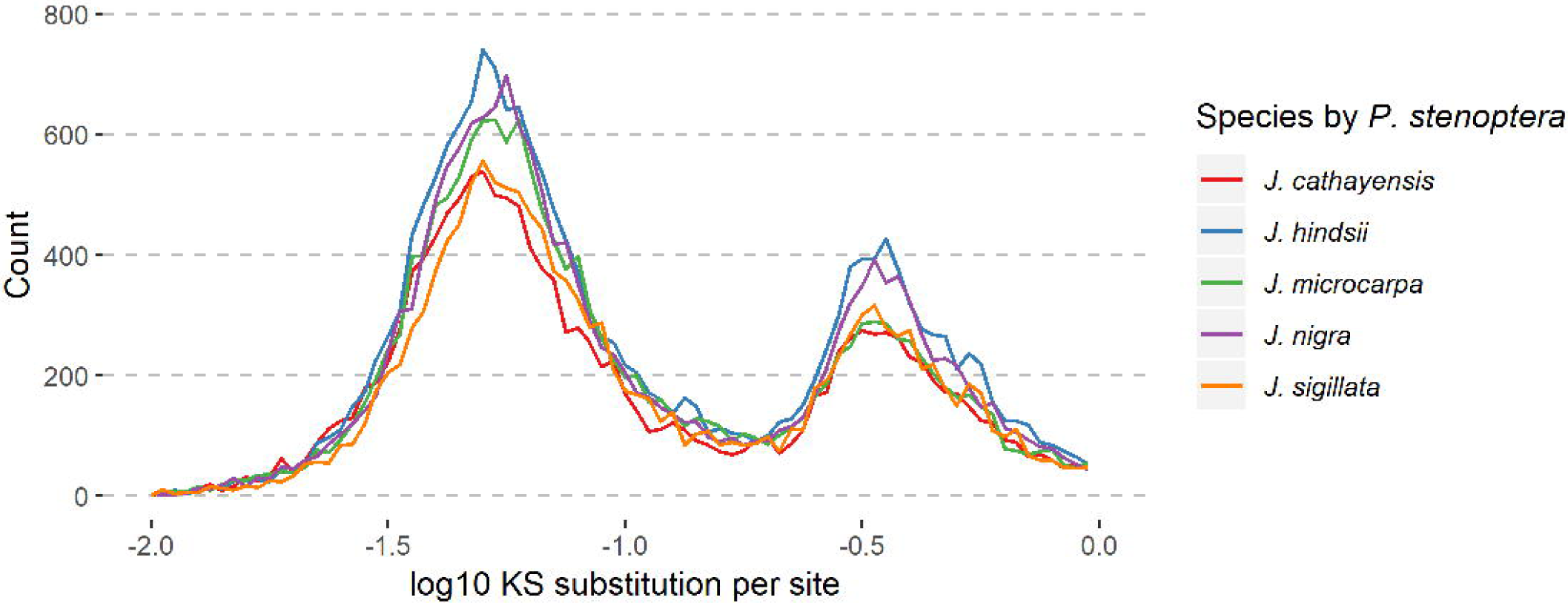
Phylogenetic tree constructed from divergence times in literature displaying numbers of expanded (blue) and contracted (red) orthogroups per terminal branch discovered using OrthoFinder/CAFÉ with the 13 species Eurosid analysis. The number of significant (P-value <0.05, Viterbi P-value <0.05) expansions and contractions at each node and leaf are shown in parentheses.

### Rhysocaryon Gene Family Evolution

At the ancestral *Rhysocaryon* node, 4 significant expansions and 2 significant contractions were discovered. Functional annotation of Juglandaceae orthogroups expanded in *J. microcarpa* revealed high incidence of transferase activity (GO:0016740) which occurred in 8 of 123 orthogroup annotations. An orthogroup annotated as ankyrin repeat-containing (OG0000093) was significantly expanded in both *J. microcarpa* (22 genes) and *J. sigillata* (16 genes) relative to other species (0-6 genes). Three orthogroups annotated as Kinesin-like protein KIN-4C, Phosphatidylinositol 4-kinase gamma 7 (P4KG7) and RNA-dependent RNA polymerase (OG0002363, OG0001584 and OG0013906) were expanded in *J. microcarpa* and *J. nigra* relative to other annotated species. An activating signal cointegrator orthogroup (OG0022144) with a zinc finger-C2HC5 (Pfam:PF06221) domain was expanded (+13) in *J. microcarpa*. OG0000386, annotated as topless-related protein 1 (TPR1) was also expanded (+6). Three orthogroups annotated as “wall-associated receptor kinase-like” (OG0000502, OG0000046 and OG0000685) lacked gene models from both *J. hindsii* and *J. microcarpa*. OG0000685 also lacked *J. sigillata* gene models. A Heat Shock Cognate 70 kDa (HSC70) orthogroup (OG0000060) was expanded in both *J. hindsii* (23 genes) and *J. microcarpa* (30 genes) relative to all species outside of *Rhysocaryon* (1-2 genes) and unexpectedly lacked gene models from *J. nigra*. Similarly, SAPK10-like serine/threonine kinase orthogroup (OG0001146) was also expanded in both *J. hindsii* (8 genes) and *J. microcarpa* (7 genes) relative to other species (0-4 genes) and lacked *J. nigra* gene models.

The large Juglandaceae callose synthase 3-like orthogroup (OG0000004) is absent in *J. nigra* and highly contracted in *J. microcarpa* and *J. cathayensis* (7 genes) relative to other species (32-36 genes). Four cyclic nucleotide-gated ion channel orthogroups involved in Plant-pathogen interaction (KEGG:04626), are lost or highly contracted in *J. nigra*: OG0000038 (−7), OG0000567 (−3), OG0000177 (−4), OG0000603 (−2). Juglandaceae REDUCED WALL ACETYLATION 2 (RWA2) (OG0000145), putative disease resistance protein (OG0000022) and receptor-like protein kinase FERONIA-like (OG0000471) orthogroups lacked *J. hindsii* gene models despite being represented by every other species.

### Dioscaryon Gene Family Evolution

At the ancestral *Dioscaryon* node, 5 significant expansions and 8 significant contractions were discovered. Annotated gene family expansions specific to *Dioscaryon* include (+3) ABC transporter B family orthogroup (OG0000286), and (+2) Ethanolamine-phosphate (OG0001347) orthogroups. Juglandaceae cationic peroxidase (CEVI16) orthogroup (OG0000173) related to Phenylpropanoid biosynthesis (GO:0009699) is contracted in *Juglans* sect. *Dioscaryon*. Probable reticuline oxidase families (OG0000562, OG0000531) annotated as containing BBE (Pfam:PF08031) and FAD binding 4 (Pfam:PF1565) lack *Dioscaryon* gene models while all non-*Dioscaryon* species contribute at least 3 gene copies in each orthogroup. *Dioscaryon* gene models were absent in nodulin-like orthogroup (OG0000206) (−3 genes). Contractions in F-box protein orthoroup (OG0000054) and Oxygen-evolving enhancer protein 2 (OG0000122) were also observed (−4 and −3 genes, respectively). One SWIM zinc finger orthogroup (OG0000510) lacked gene models in *J. regia, J. sigillata* and *J. microcarpa*. Another orthogroup (OG0000266) annotated as SWIM zinc finger appeared to also be absent in *J. regia, J. sigillata* and *J. microcarpa*, but a *J. sigillata* ortholog was discovered as a loss through the absence of protein to genome alignment.

Gene family expansions in *J. regia* include (+4) 26s proteasome regulatory subunit (OG0019963), (+3) thaumatin-like protein (OG0000263), (+4) STOMATAL CYTOKINESIS DEFECTIVE 1-like (OG0001205), (+3) mitogen-activated protein kinase kinase kinase (OG0012715), (+4) Hydroxyproline O-galactosyltransferase GALT6 (OG0004422), (+10) tubulin beta-6 chain (OG0000238). Expanded orthogroups in *J. sigillata* include (+11) endoribonuclease dicer (OG0000131).

### Gene Family Evolution Enrichment

EggNOG gene descriptions of rapidly evolving gene families were examined to infer the major functional categories of rapidly expanding and contracting gene families across Juglandaceae. Of the 333 instances of gene family contraction calculated across the Juglandaceae, the most frequent GO molecular function terms, included: 26 transferase activity (GO:0016740), 9 lyase activity (GO:0016829), and 9 cyclase activity (GO:0009975) families. High occurrence EggNOG-derived gene family descriptions of contracting orthogroups included 25 that contained “resistance”, 54 containing “kinase”, 8 “cytochrome P450” and 7 “channel”. For the 428 instances of gene family expansion, the most frequent molecular function annotations were 29 transferase activity (GO:0016740), 8 transmembrane transporter activity (GO:0022857) and 7 heterocyclic compound binding (GO:1901363). High occurrence EggNOG descriptions of expanding orthogroups include 51 containing “kinase”, 19 that contained “resistance” 17 that contained “synthase”. Comparisons of annotated rapidly evolving gene families among Juglandaceae species did reveal disproportionate gains and losses. *J. microcarpa*, for example has 7 instances of expansion in “synthase” orthogroups while *J. sigillata* has 0 and *J. hindsii* demonstrates contraction of 10 “kinase” orthogroups, while only 2 such contractions were calculated in *J. cathayensis* (0 at the preceding node shared with *Dioscaryon*). These divergent patterns of gene family evolution underline the importance of having comprehensive genetic resources for multiple species within a single clade. The six *Juglans* genome annotations provide an immediate reference for one another and construct a genetic background for the genus.

Of the 153 significant gene family size changes in *J. microcarpa*, 131 represent expansions. The changes in other *Juglans* species are more evenly distributed between expansions and contractions. The inflated number of significant expansions in *J. microcarpa* likely reflects uncollapsed heterozygosity left behind by the genome assembly process, especially given the unexpectedly large size of the *J. microcarpa* assembly (Table 1). A similar, but less pronounced pattern is observed in *J. cathayensis*.

### Selection Analysis

The likelihoods of one-ratio (null), nearly neutral (NN) and positive selection (PS) models were compared (Table S7). Of the 15 gene families that were tested, the nearly neutral model fit the data significantly better than the null for 2 orthogroups and the positive selection model for 10. Of these, 6 orthogroups (OG0000038, OG0000567, OG0000603, OG0001146, OG0001205, OG0001222) were found to be under positive selection across the selected sequences. OG0000038 (PS ω = 1.67), OG0000567 (PS ω = 1.76) and OG0000603 (PS ω = 1.84) were annotated as cyclic nucleotide-gated ion channel proteins, OG0001146 (PS ω = 1.24) as a serine threonine-protein kinase, OG0001205 (PS ω = 9.96) annotated as STOMATAL CYTOKINESIS DEFECTIVE 1-like and OG0001222 (PS ω = 2.42) as Chitinase-3.

### Divergence Estimates

We estimated the distribution of nucleotide substitution rates at silent codon positions between each of the *Juglans* genomes studied and the outgroup *Pterocarya stenoptera*. For each pairwise analysis, we observed similar bimodal distributions of synonymous substitution rates (Ks) between syntenic blocks of genes (Figure 4A). For these syntenic blocks of genes, a whole genome duplication event would give rise to such a bimodal distribution in time to the most recent common ancestor. For each species pair, we thus estimated the two modes of the distribution (Table S8). The estimates for the higher mode ranged from a low of Ks = 0.356 to a high of Ks = 0.364 with an average value of Ks = 0.361. The lower mode ranged from a low of Ks = 0.050 to a high of Ks = 0.054 with an average value of Ks = 0.053. While the non-synonymous substitution rates (Kn) between syntenic blocks of genes were much lower, the distributions were also bi-modal in appearance (Figure 4B)

**Figure 4:**
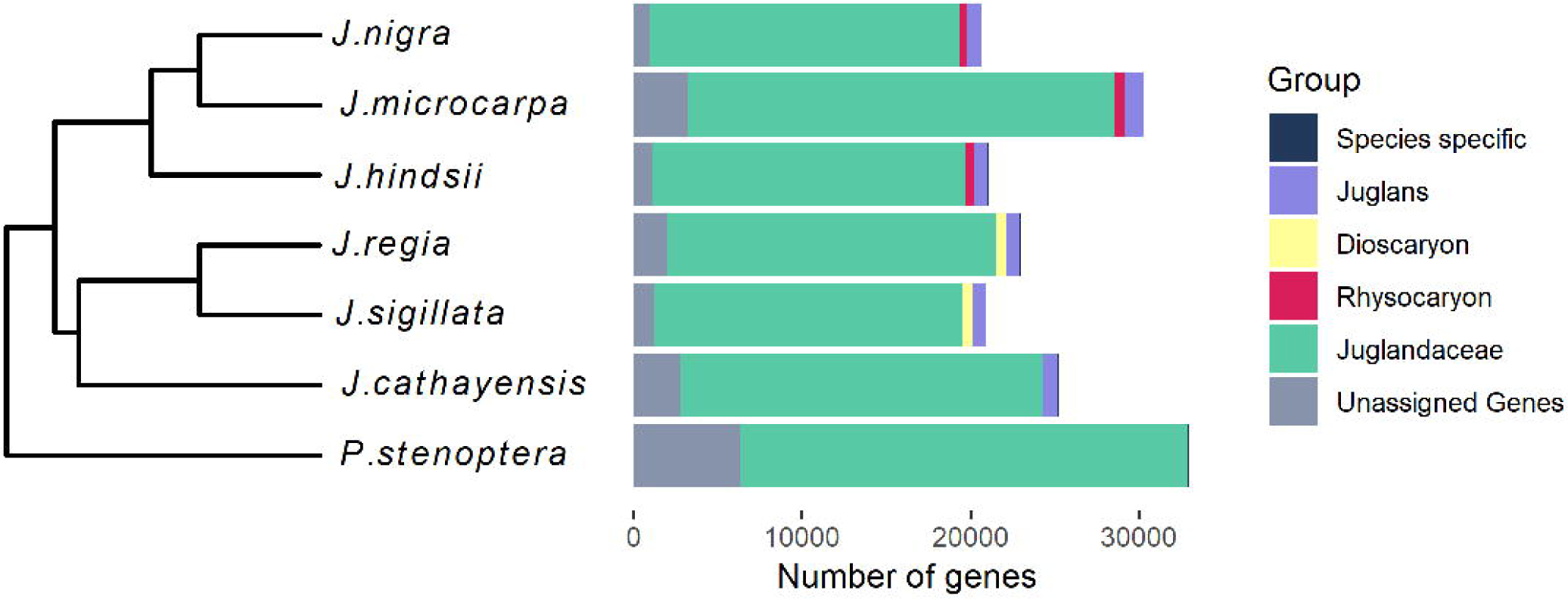
Histograms of substitution rates for coding genes determined by SynMAP to be syntenic between *Juglans hindsii* and *Pterocarya stenoptera*. Two peaks are visible in both the non-synonymous (A) and synonymous (B) distributions. In both cases the highlighted righthand peak represents the older WGD. Table S8 summarizes the distributions for all annotated *Juglans* genomes described here against *P. stenoptera*.

The annotation of the genome of *Quercus robur* (oakgenome.fr) allowed us to perform the same analysis with a species whose common ancestor predates the whole genome duplication event common to the Juglandaceae. We chose the genomes of *J. regia* and *P. stenoptera* as the best representatives of their genera. In both cases, while the histogram was much sparser due to the additional divergence, a single prominent peak was observed. For *J. regia* against *Q. robur*, it was observed at a value of Ks = 0.49 and for *P. stenoptera* against *Q. robur*, it was observed at a value of Ks = 0.53. These divergence estimates are greater than all values estimated in the *Juglans*-*Pterocarya* comparisons.

## DISCUSSION

In this study, we utilized a comprehensive *J. regia* transcriptome dataset to produce high-quality genome annotations of six recently assembled species within *Juglans* and a single member of the sister genus, *Pterocarya*. The gene model set completeness as measured by BUSCO suggests our annotation pipeline is suitable for comprehensive capture of protein-coding genes. It is still expected that limitations of single species RNA-Seq as the training input introduced some bias in the annotations for the other Juglandaceae. Although the gene prediction software, BRAKER2 seems to return far fewer false positive gene models than alternative applications, the process of removing the extraneous models remains essential to producing genome annotations that can be leveraged by the community. Still, complex plant genomes, especially those derived from short read dominant assemblies, remain challenging to annotate and existing pipelines typically introduce errors and false positives (Van Bel et al. 2019). The gene model filtration steps presented here handled multi-exonic and mono-exonic genes separately and examined both structural and functional qualities of models to permit only those of the highest confidence. This phylogenetically comprehensive set of diploid genome annotations represents an invaluable resource for comparative genomics studies within *Juglans* and for other clades (Tuskan *et al*. 2018).

The species annotated in this study represent each of the three sections of *Juglans* (*Cardiocaryon, Juglans* (syn. *Dioscaryon*) and *Rhysocaryon*) and represent fully the diversity in the genus. These annotations will serve as a platform for identifying genetic underpinnings of high-value agricultural characteristics such as drought tolerance and disease resistance that are scattered across the various species (Bernard *et al*. 2018). Moreover, they have the potential to add a new dimension to the ongoing medicinal natural products search within *Juglans* (Yao *et al*. 2012; Xu *et al*. 2013; Kim *et al*. 2018).

Because this dataset is representative of the diversity in *Juglans*, it allows for exceptional resolution of patterns in gene family evolution. Multiple samples within sect. *Rhysocaryon* and sect. *Dioscaryon* increase confidence that observed patterns across those genomes are true and not artifacts of technological and biological challenges.

### Challenges in Assessing Gene Family Evolution

Given the nature of short read assemblies, the possibility of an assembly or annotation error resulting in an incorrect consensus and falsely ‘pseudogenizing’ a gene model is non-zero. These errors, especially in small gene families, could be interpreted as significant contractions in the CAFE analysis. The weighty consequence of this effect on interpreting gene family evolution underscores the importance of deep sequencing for comparative studies, and as the shift towards long read sequencing progresses, adherence to best base-calling and polishing practices.

The risk of introducing false positive expansions is most prominent in the genome assembly phase. High heterozygosity in parts of a genome make the recovery of both haplotypes (for diploids) difficult for those regions. In final assemblies the haplotypes are often reported in separate contigs. Any gene models prevailing in these regions will falsely occur in duplicate within the annotation if the haplotigs are not recognized. The *J. microcarpa, P. stenoptera*, and to a lesser extent, *J. cathayensis* genomes exhibited these patterns by showing high duplication rates in BUSCO analyses (Table S4), inflated numbers of gene models (Table S2), and larger than expected genome sizes (Table 1). The evidence for uncollapsed heterozygosity in these genomes was reinforced by the absence of an additional peak representing taxa-specific duplications in the Ks distributions. Computational tools have been developed to address the challenges of resolving heterozygous region but are most effective when applied to long-read (or hybrid) assemblies (Chin et al. 2016).

The vastly reduced cost of sequencing over the past several years has enabled genus-level analysis of whole genome diversity, a scale at which it becomes tractable to assess patterns and significance of changes in gene CNV and other structural variation. Given the newness of this capability, a sharp increase in sequencing projects capable of resolving CNV should be expected. However, there are still only a few studies that have established the phenotypic and fitness consequences of CNV (Cook *et al*. 2012, Würschum *et al*. 2018) and even fewer that involve full-genome assessments (Prunier *et al*. 2018).

Convergent shifts in copy number under strong selective pressure for glyphosate resistance were reported for the *EPSPS* gene in eight weedy species (Patterson *et al*. 2018). This finding is notable because it points towards modulated gene expression levels through CNV as a potential source of rapid adaptation on short timescales. These types of structural variations most often occur in genomic regions called CNV hotspots, which are enriched for low-copy repeats (LCRs) (Hastings *et al*. 2009). In a genome-wide survey, distinguishing between an ancestral event and parallel evolution would require attention to the entire duplicated genomic region in each taxon. These investigations lend a greater importance to the production of near chromosome-level assemblies because poor contiguity obscures the ability to resolve structural variants.

A recent pangenome study in *Poplar* showed that intraspecific CNV occurred across each of the three genomes sequenced from hybridizable species (Pinosio *et al. 2016*). This and similar studies suggest that a single genome assembly from a single locality is likely not representative of the copy number diversity that exists within the sampled population (Hirsch et al. 2014; Golicz et al. 2016; Gordon et al. 2017; Zhao et al. 2018).

By this notion, the following observations are in no way confirmatory without additional sources of evidence. Although this dataset does not resolve interspecific diversity, it is still representative of the diversity in Juglans, and allows for exceptional resolution of patterns in gene family evolution. Multiple samples within sect. *Rhysocaryon* and sect. *Dioscaryon*, and careful attention to informatic strategies, increases confidence that the observed patterns across these genomes are true and not artifacts.

### Disease Resistance

#### Losses in Dioscaryon

The absence of *Dioscaryon* gene models in the reticuline oxidase (berberine bridge enzyme, BBE) annotated orthogroups shows a contraction before their divergence 22 MYA (Stevens *et al*. 2018). Enzymes in this family have been shown to contribute to alkaloid production (Fujii *et al*. 2007) in California poppy (*Eschscholzia californica*) and have been implicated in monolignol metabolism. Extreme (400-fold) upregulation of enzymes in this family has been observed during pathogen attack and osmotic stress in *Arabidopsis* (Daniel *et al*. 2015). Recent work in *Arabidopsis* demonstrated the function of one BBE-like enzyme in oxidizing oligogalacturonides (OGs) and thereby diminishing their elicitor activity (Benedetti *et al*. 2018). It is likely that the loss of the BBE gene family in *J. regia* and *J. sigillata* occurred in the *Dioscaryon* ancestor but that does not eliminate the possibility that these species were favored and therefore selected for their potentially tamed secondary metabolite profiles. Until recently, chemical analyses in *Juglans* have been limited to observational studies and comparisons of different cultivars within a species (Vu *et al*. 2018; Vu *et al*. 2019). Additional studies contrasting metabolomic profiles of domesticated species with their wild relatives will offer valuable insight into tree domestication, especially when paired with genome annotations.

The wall-associated kinases (WAKs) are a family of transmembrane receptor-like proteins that bind pectin in the extracellular matrix (ECM) (Wagner and Kohorn, 2001). They are necessary for cell expansion in Arabidopsis seedlings, but when bound to OGs also function in defense response through Enhanced disease susceptibility 1 (EDS1) and Phytoalexin deficient 4 (PAD4) dependent activation of MPK6-dependent pathway (Kohorn et al. 2009; Brutus et al. 2010; Kohorn et al. 2014; Davidsson et al. 2017). Recent studies of WAKs have shed light on their role in plant response to abiotic stressors (Marakli and Gozukirmizi. 2018, Xia et al. 2018) but many WAK family genes remain without functional characterization. Because of this, the parallel contraction and loss of J. hindsii and J. sigillata genes from multiple WAK annotated orthogroups is difficult to speculate on. A more elaborate depiction of the WAK gene family will certainly shed light on the significance of these losses. It is interesting to note, however, that two gene families (WAK and BBE) which have members known to interact with OGs are both contracted in J. sigillata. These losses suggest a significant shift in J. sigillata effector-triggered immunity.

#### Losses and contractions in J. hindsii

In California, the cultivation of *J. regia* is most commonly facilitated using Paradox rootstock (*J. hindsii* 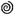 *× J. regia* 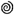), which is valued for its resistance to soil-borne pathogens (Browne *et al*. 2015, Potter *et al*. 2002). Despite higher resistance to several diseases, Paradox rootstock remain susceptible to *Armillaria* root rot, which is caused by a basidiomycete, *A. mellea* in California (Baumgartner *et al*. 2013). The impact of this disease is worsened by the lack of post-infection controls. Accordingly, discovering resistant Paradox hybrids has been the focus of some research, but has achieved limited success relative to the levels of *Armillaria* resistance reported in *J. hindsii* (Drakulic *et al*. 2017). Full genome annotations for these *Juglans* and others might be able to impart clues about the genetic distinctions that contribute to these agriculturally interesting phenotypes.

For comparison, interactions between *Arabidopsis* and the fungal pathogen, *Fusarium oxysporum* are intensely studied as a model for plant fungal diseases. In this system, *F. oxysporum* infections are potentiated by the alkalinization of soil around host root tissue caused by plant RALF-triggered alkalinization response to pathogen secreted peptides, RALFs (Rapid Alkalinization Factors) homologs (Masachis *et al*. 2016). These fungal peptides target various members of transmembrane receptor-like kinases encoded by the plant *Catharanthus roseus* Receptor-like Kinase (CrRLK1L) gene family. There are 17 reported CrRLK1L protein orthologs in *Arabidopsis* that have been implicated in a variety of processes including immunity signaling, abiotic stress response and cell wall dynamics (Kessler *et al*. 2010, Richter *et al*. 2017, Guo *et al*. 2018, Richter *et al*. 2018). Several Basidiomycete genomes have been reported to encode RALF homologs, making it plausible that *Armillaria* is among the fungi that utilize RALF-homolog effectors in infection.

The current literature suggests a central role for CrRLK1L with respect to *F. oxysporum* resistance. It is possible that the reduction or absence of the CrRLK1L orthologs (annotated as FERONIA) in *J. hindsii* is at least partially responsible for its resistance to *A. mellea* infection. The absence of *J. hindsii* models across five orthogroups annotated as receptor-like protein kinase FERONIA-like in the Juglandaceae comparison (OG0000687, OG0000471, OG0022392, OG0009045, OG0013911) including two for which every other species is represented (OG0000687, OG0000471) warrants further investigation. If the gene family is lost in *J. hindsii*, discovering any compensatory mechanisms that might maintain the integrity of CrRLK1L-involved pathways could have application in engineering fungus-resistant plants.

Like the observation that *Arabidopsis* FERONIA knockouts are more resistant to *Fusarium* infection (Masachis *et al*. 2016), the loss of function *Arabidopsis* mutations in RWA2 (reduced wall acetylation-2) led to increased resistance against the Ascomycete pathogen, *Botrytis cinerea*, the causal agent of grey mold (Manabe *et al*. 2011). *B. cinerea* belongs to a family of fungi, Botryosphaeriaceae, several of which are known to infect the nuts of *J. regia* and related species (Moral et al. 2010). The Juglandaceae RWA2 orthogroup (OG0000145) was missing *J. hindsii* gene models. RWA2 is involved in secondary cell wall synthesis and is regulated by SND1 (secondary wall-associated NAC domain protein 1) (Lee *et al*. 2011). No experiments to date have assessed the susceptibility of various *Juglans* species to *B. cinerea*, but it would be interesting to examine resistance to the pathogen in a species without RWA2. The observed loss of Chitinase-3 (Cht3) (OG0001222) in *J. hindsii* is consistent with the loss of FERONIA and RWA2. Cht3 (along with glucanase and thaumatin-like protein) are aspects of plant response to fungal invasion (Singh *et al*. 2012) and was found to be under positive selection in the additional *Juglans* species (Table S7). If the loss of FERONIA and RWA2 do correspond to a weakened compatible host signature, the decreased incidence of fungal infection would render such defense responses inessential.

### Mutation rate estimates and evidence of WGD

Testing for WGD supported the hypothesis of a Juglandoid duplication. We observed similar bimodal distributions of Ks values among syntenic blocks of genes in each of the *Juglans-Pterocarya* pairwise analyses (Figure 4A). The bimodal distribution can be attributed to a mixture of estimates from two distinct lineages; comparisons between orthologous genes; and comparisons between more distant paralogous genes arising from the whole genome duplication. Bimodal distributions for all *Juglans-Pterocarya* pairwise comparisons are consistent with the WGD occurring prior to the radiation of Juglans (Luo et al. 2015; Zhu et al. 2019). As additional confirmation, the most prominent feature in the same analysis against the annotated genome of *Quercus robur* is a peak at divergence values greater than those estimated for the WGD (Table S5).

Using the larger mode for each of the five distributions, we can estimate the nucleotide substitution rate using the method of Zhu *et al*. (2019), for comparison. Using 66 MYA as the assumed date of the WGD from Zhu *et al*. (2019), we obtain a synonymous mutation rate of 2.7×10-9. This rate is higher than the rate of 2.3×10-9 estimated using 14 genes in Luo et al. (2015) and closer to the more recent estimate of 2.5×10-9 in Zhu *et al*. (2019) using thousands of genes in a *J*.*regia* × *J*.*microcarpa* hybrid. Our faster rate is still more consistent with the rates of other woody perennials (e.g. Palm (Gaut *et al*. 1996) and *P. trichocarpa* (Tuskan *et al*. 2006)), and still five times slower than the rate reported for *Arabidopsis* (Koch *et al*. 2000).

A surprising observation was the distance between the two modes. We assume that the estimated Ks of the smaller mode represents the between species divergence. The ratios of the larger to smaller modes ranged from 6.5 to 7.3. Interpretation of fossil data (Manchester 1987) placed the initial split into *Rhysocaryon* and *Cardiocaryon* around 45 MYA, resolving around 38 MYA. Much closer to the assumed time of the WGD than our bimodal distributions of Ks would indicate under a molecular clock. The discrepancy in estimated WGD times may be due to the non-neutral nature of these substitutions and departure from a molecular clock. However, a relevant observation was recently made using a coalescent based approach. Bai *et al*. (2018) noted that convergence of effective population size indicates a much earlier beginning for the divergence among Juglans lineages. Our data could also be interpreted to support a more recent divergence of walnut lineages.

The resources and services provided by *Juglans* species are nutritionally and culturally significant. Their wood, used to construct furnishing and musical instruments, is valued among woodworkers. Ink from walnut husks was used by Leonardo da Vinci and Rembrandt. Brown dye from walnut stained fabrics was used in classical Rome, medieval Europe, Byzantium and the Ottoman Empire. The genus is elevated in poetry across the globe, including for its non-monetary benefits in Mary Oliver’s “The Black Walnut Tree” (Oliver, 1992) and nutritional properties in Tatsuji Miyoshi’s “In Praise of a Walnut” (Miyoshi, 1946). We are enthusiastic to contribute to the understanding of and appreciation for this genus by constructing these genome annotation resources.

## METHODS

### Repeat Library Generation and Softmasking

The seven assemblies, ranging in size from 600 Mb to just under 1 Gb (2n=32) were assessed for repeat content (Stevens *et al*. 2018). Scaffolds and contigs less than 3Kbp in length were removed from the assemblies prior to annotation. RepeatModeler (v1.0.8) was used to construct a repeat library through a combination of *de novo* and structural prediction tools wrapped into the pipeline (Smit and Hubley, 2008). RepeatModeler provided base annotations for the repeat elements (Table S1) and generated a consensus library that was used as input to Repeatmasker (v4.0.6) to generate softmasked genomes (Smit *et al*. 2013).

### Structural Annotation

After softmasking, a set of 19 independent *J. regia* tissue-specific libraries described in Chakraborty *et al* (2015) were aligned to the reference genomes via TopHat2 (v2.1.1) (Kim *et al*. 2013). The Illumina 85bp PE sequences were independently quality controlled for a minimum length of 45bp and a minimum Phred-scaled quality score of 35 via Sickle (v. 1.33) prior to alignment. Independent alignment files were sorted and provided to Braker2 (v2.0) which generated a hints file for semi-supervised training of the *ab initio* gene prediction package, Augustus (Stanke *et al*. 2008). Braker2 utilizes RNA-Seq reads directly to inform gene prediction and deduce the final models (Hoff *et a*l. 2016). The annotation files (GFF) produced were processed by gFACs, to filter out incomplete or improbable gene models on the basis of completion (identifiable start and stop codons) and canonical gene structure (micro-exons and micro-introns < 20bp are filtered to reduce erroneous models). The gFACs package also resolves conflicting models and reports splice site statistics as well as other basic gene structure statistics (Caballero and Wegrzyn, 2019).

### Functional Annotation

The EnTAP functional annotation package was employed to remove unlikely gene models and provide provisional functional information (Hart *et al*. 2019). Multi-exonic and mono-exonic gene models were subjected to different functional filtering pipelines that each utilized EnTAP. For multi-exonic genes, EnTAP (v 0.8.1) was provided three curated databases (NCBI’s Plant Protein (release 87), NCBI’s RefSeq Protein (release 87), and UniProtKB/Swiss-Prot) for similarity search (50% target and query coverage; Diamond E-value .00001), followed by gene family assignment via the EggNOG database and EggNOG-mapper toolbox (Jensen *et al*. 2008). Associations to gene families provided the basis for Gene Ontology term assignment, identification of protein domains (PFAM), and associated pathways (KEGG) (Finn *et al*. 2014; Ashburner *et al*. 2017). Multi-exonics were removed from the set if they had neither sequence similarity search result nor gene family assignment. Mono-exonic genes are typically over-estimated in the process of *ab initio* genome annotation. To reduce this effect, they were aligned to a custom curated database of monoexonic genes from other plant species using 80% query coverage and 80% target coverage cutoffs in an independent similarity search through EnTAP. EggNOG and PFAM were used in mono-exonic gene model filtering as they were for multi-exonic filtering. After the first round of filters, InterProScan (v5.25) was used to confirm gene family assignment and protein domains in monoexonic gene models. Gene models without InterProScan annotations were removed from the monoexonic set. For each species, the filtered multi-exonic and mono-exonic gene sets were combined and passed back to gFACs to generate a statistical profile and consistent annotation file in gene transfer format (GTF). Finally, gene models that annotated with domains specific to retroelements were further filtered from the final annotations based upon Pfam database descriptions. The entire set of filtered gene models was evaluated for completeness. BUSCO (v3.0.2) was used with default parameters and the embryophyta reference set of 1440 orthologs for this purpose (Simão *et al*. 2015). Using the output from Augustus, we used gFACs to also capture partial gene models. These were also functionally annotated used EnTAP, and then compared using the same BUSCO analysis and ortholog set.

### Gene Family Classification and Evolution

The proteins derived from the filtered genome annotations of each species were processed with OrthoFinder-Diamond (v1.1.10) to provide information about orthologous gene families. OrthoFinder is robust to incomplete models, differing gene lengths, and larger phylogenetic distances (Emms and Kelly, 2015). Gene families (orthogroups) in OrthoFinder are defined as homologous genes descended from a single gene from the last common ancestor of the species examined. It is assumed that a parental gene of each orthogroup was present in the common ancestor of the seven species investigated. Two independent runs were conducted with OrthoFinder: *Juglans* with the *Pterocarya* outgroup, and another that included these species with a set of 6 selected Eurosids (*Citrus grandis, Eucalyptus grandis, Arabidopsis thaliana, Carica papaya, Populus trichocarpa* and *Quercus robur*). Rates of gene family evolution were calculated for each orthogroup using the stochastic birth and death rate modeling implemented in CAFE (v4.1) (De Bie *et al*. 2006). Species trees were constructed by applying estimated divergence times from literature detailing rosid phylogeny to the known topology (Magallón *et al*. 2014, Dong *et al*. 2017). Large variance in gene copy number between species can lead to inaccurate calculation of birth and death rate parameters, therefore large orthogroups with more than 100 gene models were removed prior to the analyses and later analyzed separately using those parameters calculated by including only orthogroups with < 100 gene models. Orthogroups represented by a single set of paralogs were also removed because they are uninformative. Rapidly evolving gene families (orthogroups) were identified using CAFE, which models the rate of gene family evolution while accounting for the uncertainty in membership that results from imperfect genome annotation. For each set, the lambda (birth and death rate) parameter was calculated uniformly across the phylogeny. Orthogroups with a large size variance among taxa were selected using a CAFE family-wide P-values <0.05. Those orthogroups with accelerated rates of evolution were selected using branch-specific Viterbi P-values <0.05. The gene-family losses described were independently confirmed using Exonerate protein2genome alignments of the longest gene in the orthogroup to the genome of the excluded species (90% similarity and score 1000) (Slater and Birney, 2005).

Functional enrichment of rapidly evolving gene families was assessed independently for each node and leaf of the Juglandaceae cladogram and across the entire set. EggNOG gene descriptions of the longest gene model from each orthogroup were compiled into a functional background. The gene model annotations from sets of orthogroups found to be either rapidly expanding or rapidly contracting at each leaf or node were compared to that background to estimate functional enrichment within the set.

### Selection Analysis

To test for positive selection in gene families of interest, the coding sequence of gene models from each orthogroup were iteratively clustered with USEARCH (v 9.0.2132) at various identities beginning at 0.95 down to a minimum of 0.7 at intervals of 0.05. Iterative clustering was terminated once a cluster with sufficient species representation (relative to the species representation of that particular orthogroup) was produced and chosen for use in selection analysis. A multiple sequence alignment of the longest gene model from each species in that cluster was produced using Clustal Omega (v 1.2.4). The multiple sequence alignments and species tree were provided to CODEML from PAML (v 4.9) to calculate ω (dN/dS), the ratio of non-synonymous to synonymous amino acid substitutions, across two models of adaptive evolution, including nearly neutral and positive selection and the corresponding likelihood values. A likelihood ratio test was used to determine the best model for each orthogroup.

### Syntelog Analysis

Genome alignment and analysis of syntenic genes was performed for each *Juglans* genome against *Pterocarya stenoptera* using a CoGE (Lyons and Freeling, 2008) SynMAP2 analysis. Genome alignment was performed using Last. Five genes were used as the minimum number of aligned pairs for DAGchainer (Haas *et al*. 2004). Synonymous (Ks) and non-synonymous (Kn) coding sequence divergence was estimated for syntenic protein coding gene pairs with CodeML (Yang, 2007).

## Data Availability

The genomic resources described here are available at NCBI under BioProject PRJNA445704 and the transcriptomic resources under BioProject PRJNA232394. These resources are also accessible from hardwoodgenomics.org and treegenesdb.org. Functional annotations, gene models and gene transfer format (gtf) files are also available on treegenesdb.org. Scripts and detailed processes used for this study are accessible on https://gitlab.com/tree-genome-annotation/Walnut_Annotation.

## Acknowledgements

This project was supported by the California Walnut Board, USDA NIFA SCRI-Award no. 2012-51181-20027 and USDA ARS CRIS project no. 5306-22000-015-00D. We thank the Institute for Systems Genomics Computational Biology Core for access to software and hardware support and Dr. Uzay Sezen at the University of Connecticut for providing enthusiasm and suggested revisions. The authors declare no conflict of interest.

## Author Contributions

DBN, CHL, AD and KAS envisioned the resource and generated the sequence data. JLW, KAS, SZ, AJT and TF outlined the comparative methodology. AJT, TF, SZ, MC and KAS analyzed the data. AJT, TF and JLW wrote the paper.

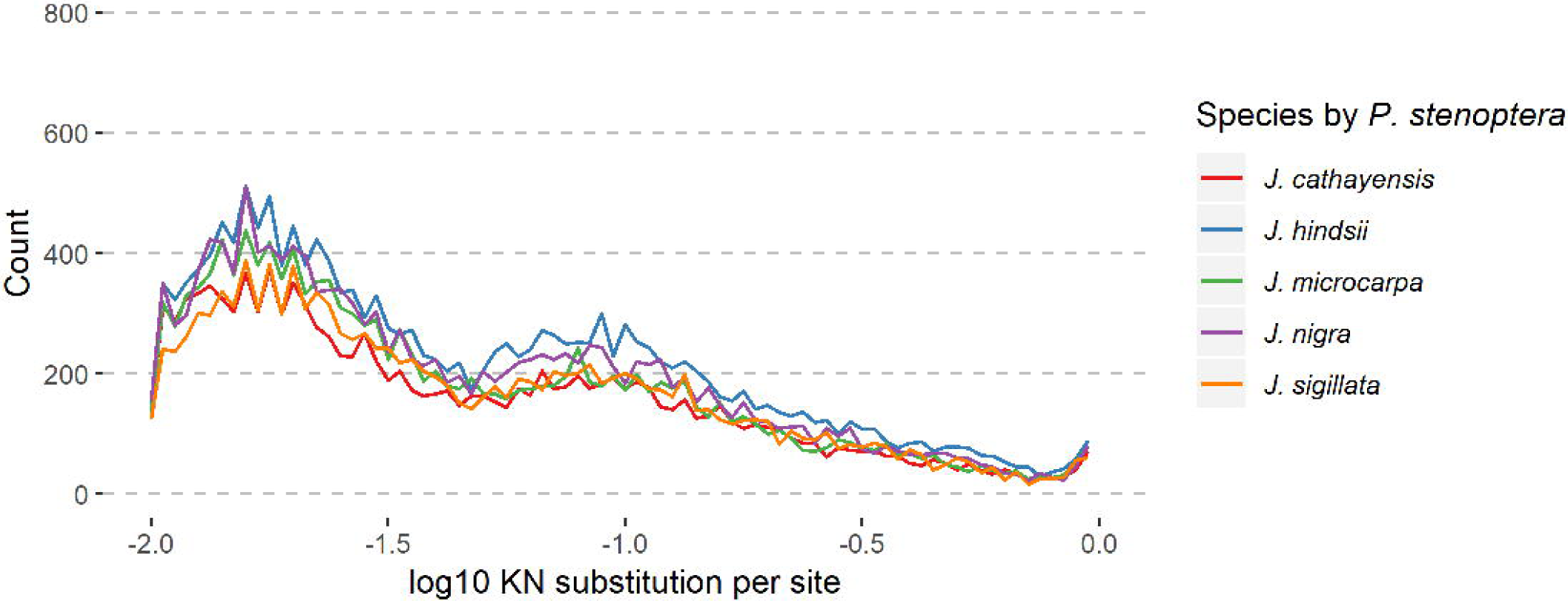

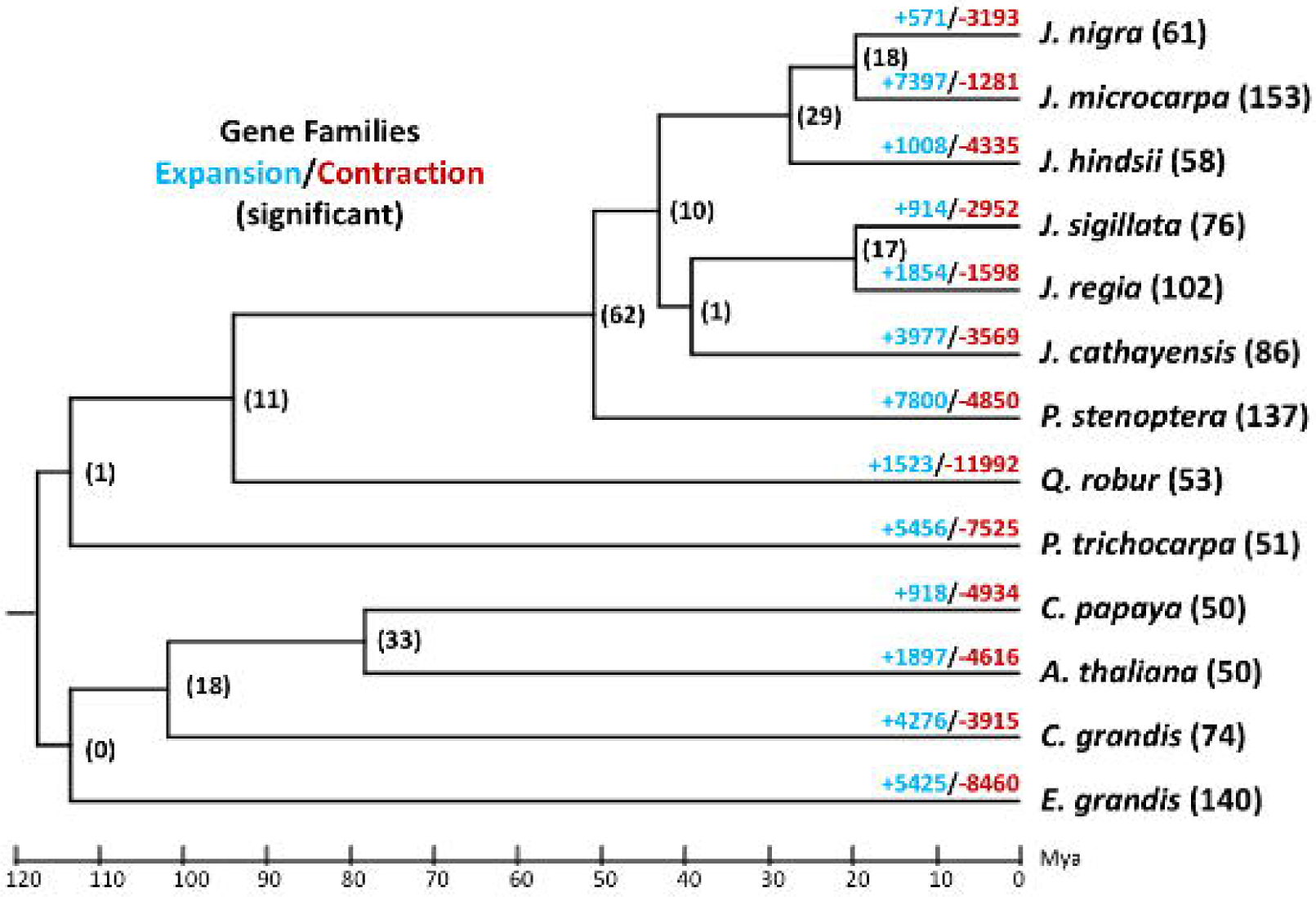

