## supplemental tables S1 - S8 for "Comparative Genomics of Six *Juglans* Species Reveals Disease-associated Gene Family Contractions"

Table S1: Repeat classification of the seven Juglandaceae genomes

|  | *Juglans hindsii* | *Juglans nigra* | *Juglans cathayensis* | *Juglans microcarpa* | *Juglans sigillata* | *Juglans regia* | *Pterocarya stenoptera* |
| --- | --- | --- | --- | --- | --- | --- | --- |
| DNA | 169,225 | 119,020 | 125,581 | 187,285 | 105,830 | 438,423 | 138,061 |
| LTR | 5,604 | 9,746 | 2,730 | 5,230 | 2,865 | 192,643 | 5,245 |
| LTR/Caulimovirus | 12,642 | 13,415 | 12,481 | 20,402 | 19,398 | 16,038 | 19,621 |
| LTR/Copia | 180,988 | 150,425 | 146,725 | 245,386 | 146,422 | 182,798 | 185,162 |
| LTR/ERV1 | 9,684 | 5,584 | 3,626 | 0 | 757 | 2,534 | 13,403 |
| LTR/ERV4 | 0 | 1,758 | 643 | 1,002 | 0 | 876 | 0 |
| LTR/ERVL | 0 | 0 | 0 | 0 | 658 | 0 | 0 |
| LTR/ERVK | 0 | 0 | 2,170 | 367 | 0 | 0 | 392 |
| LTR/Gypsy | 182,043 | 126,889 | 154,588 | 166,826 | 123,357 | 122,026 | 142,111 |
| LTR/Ngaro | 625 | 606 | 0 | 0 | 0 | 0 | 0 |
| LTR/Cassandra | 0 | 102 | 0 | 0 | 207 | 0 | 0 |
| LTR/Pao | 0 | 0 | 0 | 2,898 | 0 | 906 | 2,431 |
| nonLTR | 0 | 0 | 0 | 0 | 0 | 798 | 0 |
| LINE | 96,081 | 78,928 | 115,088 | 138,774 | 102,226 | 162,442 | 139,303 |
| SINE | 6,687 | 525 | 1,524 | 5,958 | 1,082 | 6,856 | 1,839 |
| Simple Repeats | 12,453 | 7,213 | 18,730 | 25,932 | 11,195 | 13,327 | 14,843 |
| Satellites | 0 | 6,580 | 727 | 1,415 | 10,607 | 0 | 1,570 |
| Rolling Circle (RC) | 22,532 | 19,964 | 28,955 | 28,768 | 33,051 | 27,015 | 32,959 |
| rRNA | 0 | 0 | 0 | 0 | 625 | 8,870 | 0 |
| Unknown | 479,701 | 424,502 | 506,155 | 603,858 | 495,077 | 33,226 | 539,485 |
| Total | 1,178,265 | 970,317 | 1,119,723 | 1,434,101 | 1,054,357 | 1,269,529 | 1,237,058 |

Table S2: Initial predictions of gene models provided by Braker2 and derived from RNA-Seq evidence

|  | *Juglans hindsii* | *Juglans nigra* | *Juglans cathayensis* | *Juglans microcarpa* | *Juglans sigillata* | *Juglans regia* | *Pterocarya stenoptera* |
| --- | --- | --- | --- | --- | --- | --- | --- |
| Average Gene Length | 3,158 | 2,976 | 3,003 | 2,929 | 2,993 | 3,032 | 2,985 |
| Average Number of Introns | 3.17 | 3.11 | 2.98 | 2.99 | 3.10 | 3.03 | 2.96 |
| Number of genes | 81,753 | 87,669 | 106,253 | 125,291 | 89,512 | 93,468 | 133,963 |
| Number of complete genes | 76,847 | 82,610 | 97,312 | 114,573 | 83,457 | 84,098 | 123,420 |
| Number of mono-exonic genes | 10,238 | 10,893 | 14,039 | 15,564 | 10,653 | 13,271 | 17,955 |
| Genes with either SSS (sequence similarity search) or gene family annotations | 29,382 | 29,046 | 34,857 | 43,051 | 27,596 | 31,621 | 45,808 |

Table S3: BUSCO results from the final gene models of the seven Juglandaceae species

|  | *J. cathayensis* | *J. hindsii* | *J. microcarpa* | *J. nigra* | *J. regia* | *J. sigillata* | *P. stenoptera* |
| --- | --- | --- | --- | --- | --- | --- | --- |
| Complete BUSCOs | 1,201 | 1,270 | 1,241 | 1,274 | 1,238 | 1,111 | 1,223 |
| Complete and single-copy BUSCOs | 898 | 1,101 | 726 | 1,136 | 1,069 | 975 | 662 |
| Complete and duplicated BUSCOs | 303 | 169 | 515 | 138 | 169 | 136 | 561 |
| Fragmented BUSCOs | 77 | 56 | 85 | 57 | 46 | 77 | 74 |
| Missing BUSCOs | 162 | 114 | 114 | 109 | 156 | 252 | 143 |
| Total BUSCO groups searched | 1,440 | 1,440 | 1,440 | 1,440 | 1,440 | 1,440 | 1,440 |

Table S4: BUSCO results from the gene models of the seven Juglandaceae species, including partial gene models.

|  | *J. cathayensis* | *J. hindsii* | *J. microcarpa* | *J. nigra* | *J. regia* | *J. sigillata* | *P. stenoptera* |
| --- | --- | --- | --- | --- | --- | --- | --- |
| Complete BUSCOs | 1,213 | 1,279 | 1,251 | 1,284 | 1,246 | 1,122 | 1,232 |
| Complete and single-copy BUSCOs | 903 | 1111 | 726 | 1147 | 1074 | 986 | 662 |
| Complete and duplicated BUSCOs | 310 | 168 | 525 | 137 | 172 | 136 | 570 |
| Fragmented BUSCOs | 75 | 54 | 83 | 56 | 44 | 78 | 78 |
| Missing BUSCOs | 152 | 107 | 106 | 100 | 150 | 240 | 130 |
| Total BUSCO groups searched | 1,440 | 1,440 | 1,440 | 1,440 | 1,440 | 1,440 | 1,440 |

Table S5: BUSCO results from the unannotated genome assemblies of the seven Juglandaceae species

|  | *J. cathayensis* | *J. hindsii* | *J. microcarpa* | *J. nigra* | *J. regia* | *J. sigillata* | *P. stenoptera* |
| --- | --- | --- | --- | --- | --- | --- | --- |
| Complete BUSCOs | 1,327 | 1,360 | 1,271 | 1,274 | 1,335 | 1,307 | 1,324 |
| Complete and single-copy BUSCOs | 999 | 1,192 | 726 | 1,136 | 1,138 | 1,151 | 734 |
| Complete and duplicated BUSCOs | 328 | 168 | 545 | 138 | 197 | 156 | 590 |
| Fragmented BUSCOs | 32 | 25 | 49 | 57 | 29 | 52 | 39 |
| Missing BUSCOs | 81 | 55 | 120 | 109 | 76 | 81 | 77 |
| Total BUSCO groups searched | 1,440 | 1,440 | 1,440 | 1,440 | 1,440 | 1,440 | 1,440 |

Table S6: Description of putative gene families based on two OrthoFinder runs

|  | Juglandaceae  (7 species) | Eurosid  (13 species) |
| --- | --- | --- |
| Number of genes | 234455 | 456424 |
| Number of genes in orthogroups | 216778 | 401186 |
| Number of unassigned genes | 17677 | 55238 |
| Percentage of genes in orthogroups | 92.5 | 87.9 |
| Percentage of unassigned genes | 7.5 | 12.1 |
| Number of orthogroups | 26458 | 22189 |
| Number of species-specific orthogroups | 56 | 488 |
| Number of genes in species-specific orthogroups | 161 | 3054 |
| Percentage of genes in species-specific orthogroups | 0.1 | 0.7 |
| Mean orthogroup size | 8.2 | 18.1 |
| Median orthogroup size | 8.0 | 15.0 |
| G50 (assigned genes) | 9 | 25 |
| G50 (all genes) | 8 | 23 |
| O50 (assigned genes) | 8144 | 4792 |
| O50 (all genes) | 9135 | 5965 |
| Number of orthogroups with all species present | 14429 | 6722 |
| Number of single-copy orthogroups | 2673 | 66 |

Table S7: Targeted Selection Analysis

| **Orthogroup** | **M0 (One-ratio)** | **M0 lnL** | **Nearly Neutral (NN)** | **NN lnL** | **Positive Selection (PS)** | **PS lnL** |
| --- | --- | --- | --- | --- | --- | --- |
| OG0000003 | 0.79 | -6319.06 | 0.50 | -6230.30 | 1.09 | -6183.03 |
| OG0000004 | 0.12 | -2212.84 | 0.28 | -2201.97 | 0.28 | -2201.97 |
| OG0000038 | 0.85 | -4095.00 | 0.54 | -4071.43 | 1.67 | -4044.40 |
| OG0000206 | 0.34 | -563.80 | 0.34 | -563.80 | 0.34 | -563.80 |
| OG0000238 | 0.01 | -2608.57 | 0.01 | -2582.52 | 0.01 | -2582.52 |
| OG0000286 | 0.41 | -1985.86 | 0.54 | -1978.59 | 0.78 | -1977.99 |
| OG0000386 | 0.25 | -827.56 | 0.28 | -827.29 | 0.40 | -827.14 |
| OG0000531 | 0.45 | -3002.30 | 0.34 | -2985.55 | 0.54 | -2978.35 |
| OG0000567 | 1.23 | -2824.51 | 0.54 | -2816.16 | 1.76 | -2793.89 |
| OG0000603 | 0.59 | -4236.37 | 0.45 | -4218.42 | 1.84 | -4197.15 |
| OG0001146 | 0.58 | -986.55 | 0.47 | -985.94 | 1.24 | -983.83 |
| OG0001205 | 0.96 | -1707.39 | 0.62 | -1704.55 | 9.96 | -1692.32 |
| OG0001222 | 1.22 | -3267.09 | 0.55 | -3238.07 | 2.42 | -3193.60 |
| OG0013906 | 0.41 | -1444.56 | 0.41 | -1444.56 | 0.41 | -1444.56 |
| OG0022144 | 0.16 | -980.45 | 0.18 | -979.92 | 0.18 | -979.92 |

Table S8:

|  | *J. hindsii* x  *P. stenoptera* | *J. nigra* x  *P. stenoptera* | *J. cathayensis* x *P. stenoptera* | *J. microcarpa* x  *P. stenoptera* | *J. sigillata* x  *P. stenoptera* |
| --- | --- | --- | --- | --- | --- |
| Synonymous (Ks) | | | | | |
| Lower mode | 0.053 | 0.053 | 0.050 | 0.053 | 0.054 |
| Higher mode | 0.361 | 0.361 | 0.361 | 0.364 | 0.356 |
| ratio | 6.9 | 6.8 | 7.3 | 6.8 | 6.5 |
| Non-synonymous (Kn) | | | | | |
| mode 1 | 0.017 | 0.017 | 0.016 | 0.017 | 0.018 |
| mode 2 | 0.084 | 0.078 | 0.083 | 0.083 | 0.082 |
| ratio | 5.0 | 4.7 | 5.3 | 5.0 | 4.6 |
